# Optical Estimation of Absolute Membrane Potential Using One- and Two-Photon Fluorescence Lifetime Imaging Microscopy

**DOI:** 10.1101/2021.02.16.431491

**Authors:** Julia R. Lazzari-Dean, Evan W. Miller

**Affiliations:** Department of Chemistry, University of California, Berkeley; Department of Molecular and Cell Biology, and University of California, Berkeley; Helen Wills Neuroscience Institute, University of California, Berkeley

## Abstract

**Background:** Membrane potential (V_mem_) exerts physiological influence across a wide range of time and space scales. To study V_mem_ in these diverse contexts, it is essential to accurately record absolute values of V_mem_, rather than solely relative measurements.

**Materials & Methods:** We use fluorescence lifetime imaging of a small molecule voltage sensitive dye (VF2.1.Cl) to estimate mV values of absolute membrane potential.

**Results:** We test the consistency of VF2.1.Cl lifetime measurements performed on different single photon counting instruments and find that they are in striking agreement (differences of <0.5 ps/mV in the slope and <50 ps in the y-intercept). We also demonstrate that VF2.1.Cl lifetime reports absolute V_mem_ under two-photon (2P) illumination with better than 20 mV of V_mem_ resolution, a nearly 10-fold improvement over other lifetime-based methods.

**Conclusions:** We demonstrate that VF-FLIM is a robust and portable metric for V_mem_ across imaging platforms and under both one-photon and two-photon illumination. This work is a critical foundation for application of VF-FLIM to record absolute membrane potential signals in thick tissue.

## Introduction

Membrane potential (V_mem_), or a voltage arising from ionic concentration gradients across semi-permeable membranes, plays diverse and important roles in biological systems. Of particular note is the range of time and length scales over which V_mem_ can influence cellular and organismal physiology.^1–3^ On short timescales, V_mem_ changes control neuronal communication and cardiomyocyte contraction. However, all cells, even non-electrically excitable cells, maintain a transmembrane potential, spending an estimated 10-50% of their cellular ATP budget to pump ions in opposing directions.^4^ On longer timescales, V_mem_ displays diverse patterns, including responding to growth factor signals,^5,6^ oscillating throughout the cell cycle,^7^ and marking developmental boundaries in tissue.^3,8^ Further, these voltage signals can be compartmentalized into small areas such as dendritic spines^9^ or delocalized over larger tissues.^10^

Accurate and non-invasive V_mem_ recording techniques are required to document and understand the many roles of V_mem_. On one hand, patch-clamp electrophysiology, while highly accurate, is damaging to cells and low throughput. On the other hand, recently developed optical V_mem_ recording techniques using voltage sensitive proteins or small molecules have enabled higher throughput and less invasive recordings, at the cost of voltage resolution. These optical strategies have been particularly successful in reporting on fast changes in V_mem_ in electrically excitable systems, such as neuron or cardiomyocyte action potentials, but they cannot generally report on an absolute V_mem_ (in, e.g., millivolts). Additionally, the ability to monitor changes in V_mem_ via two-photon excitation (2P)^11^ would aid in monitoring V_mem_ changes and measuring absolute V_mem_ in model systems of increasing complexity and size.

To address this need, we recently reported VF-FLIM,^6^ a technique for monitoring absolute V_mem_ across the plasma membrane with the fluorescence lifetime (τ_fl_) of the VoltageFluor dye VF2.1.Cl.^12^ Fluorescence lifetime is a measure of how long a fluorophore stays in the excited state before emitting a photon. Because it is an intrinsic property of a dye in its environment, fluorescence lifetime can be used to quantitatively make measurements in cells.^13^ In our previously published work, we demonstrated that the photoinduced electron transfer (PeT)-based mechanism of VF2.1.Cl^12,14^ leads to a linear response of lifetime with respect to V_mem_.

In this paper, we expand upon VF-FLIM, showing that our measurements are robust across instruments (two additional TCSPC-FLIM systems) and that VF-FLIM can be used under one-photon or two-photon illumination. We also describe some considerations for image analysis and provide a code package that we use for analyzing FLIM data of fluorescence associated with membranes (FLIM <u>f</u>luorescence <u>l</u>ifetime <u>a</u>nalysis <u>m</u>odule, or FLIM-FLAM). To our knowledge, this is the first demonstration of two-photon illuminated absolute V_mem_ recordings with voltage resolution better than 20 mV.

## Materials and Methods

### Materials

VF2.1.Cl was synthesized in-house according to the published synthesis.^12^ VF2.1.Cl was stored as a 1000x (100 µM) stock in DMSO at −20°C, and stocks were checked by LC-MS every few months to confirm no decomposition had occurred. All chemicals were obtained from either Sigma-Aldrich or Thermo Fisher Scientific.

### Cell Culture

HEK293T cells were obtained from the UC Berkeley Cell Culture Facility and were verified by STR profiling. Cells were maintained in a humidified 37°C incubator with 5% CO_2_ and were discarded after thirty passages. Cells were maintained in Dulbecco’s Modified Eagle Medium (DMEM) supplemented with 4.5 g/L glucose, 2 mM GlutaMAX, and 10% fetal bovine serum (FBS). Cells were dissociated with 0.05% trypsin-EDTA for passaging and preparation of microscopy samples. FBS was purchased from Seradigm; all other media and supplements were purchased from Gibco (Thermo Fisher Scientific).

14 to 24 hours before electrophysiology recordings, HEK293T cells were dissociated and plated at a density of 21,000 cells/cm^2^ in low glucose DMEM (1 g/L glucose, 2 mM GlutaMAX, 10% FBS, 1 mM sodium pyruvate) on poly-D-lysine coated 25 mm coverslips (Electron Microscopy Sciences) in a 6 well tissue culture plate (Corning). Coverslips were prepared by acid wash in 1 M HCl for 2-5 hours, followed by three overnight washes in 100% ethanol and three overnight washes in MilliQ purified water. Coverslips were sterilized by heating to 150°C for 2-5 hours. Before addition of cells, coverslips were incubated with poly-D-lysine (Sigma-Aldrich, made as a 0.1 mg/mL solution in phosphate-buffered saline with 10 mM Na_3_BO_4_) for 1-24 hours in a humidified 37°C incubator and washed twice with sterile MilliQ purified water and twice with dPBS.

### Microscopy Sample Preparation

Cells were loaded with 100 nM VF2.1.Cl for 20 minutes in a humidified 37°C incubator with 5% CO_2_ in imaging buffer (IB; pH 7.25; 290 mOsmol/L; composition in mM: NaCl 139.5, KCl 5.33, CaCl_2_ 1.26, MgCl_2_ 0.49, KH_2_PO_4_ 0.44, MgSO_4_ 0.41, Na_2_HPO_4_ 0.34, HEPES 10, D-glucose 5.56). Cells were washed once in IB and transferred to fresh IB for electrophysiology.

### Whole Cell Patch Clamp Electrophysiology

Electrodes were pulled from glass capillaries with filament (Sutter Instruments) with a P-97 pipette puller (Sutter Instruments) to resistances of 4-7 MΩ. Electrodes were filled with a K-gluconate internal solution (pH 7.25; 285 mOsmol/L, composition in mM: 125 potassium gluconate, 10 KCl, 5 NaCl, 1 EGTA, 10 HEPES, 2 ATP sodium salt, 0.3 GTP sodium salt). EGTA (tetraacid form) was prepared as a stock solution in 1 M KOH before addition to the internal solution. Voltage steps were corrected for the calculated liquid junction (pClamp software package, Molecular Devices) between IB and the K-gluconate internal.^15^

Electrodes were position with an MP-225 micromanipulator (Sutter Instruments) to obtain a gigaseal prior to breaking into the whole cell configuration. Recordings were sampled at a rate of > 10 kHz using an Axopatch 200B amplifier, filtered with a 5 kHz low-pass Bessel filter, and digitized with a DigiData 1440A (Molecular Devices). Only recordings that maintained a 30:1 ratio of membrane resistance R_m_ to access resistance R_a_ were used for analysis. Pipette capacitance was corrected with the fast magnitude knob only; series resistance compensation was not performed. Voltage steps of −80, −40, 0, and +40 mV were applied in random order, followed by a voltage step to +80 mV.

### Nikon A1R-HD25 on Ti2E with Becker & Hickl FLIM

Measurements were performed on a Nikon A1R-HD25 confocal on a Ti2E base. Excitation was provided by a 488 nm pulsed diode laser (repetition rate 50 MHz) and directed to the sample with a line pass dichroic. Emission was collected through a 40x oil immersion objective immersed in Immersol 518F (Zeiss) and was filtered through an additional emission filter (488 nm long pass) before it reached the detector. Individual photons were detected with a hybrid detector (HPM-100-40, Becker and Hickl) and converted to photon arrival times with an SPC-150 photon counting card (Becker and Hickl). Acquisition times of 30 seconds were used at each V_mem_ step. A confocal pinhole of ~2-3 Airy units was used to increase photon counts at the expense of optical sectioning. Fluorescence lifetime images were acquired in Nikon Elements and transferred to SPCImage for fitting of exponential decays. VF2.1.Cl fluorescence lifetime was modeled as a biexponential decay at each pixel of the resulting FLIM image. Data were processed with the following fit parameters fixed: shift 0, offset 0. Images were binned by a factor of 1 prior to analysis (as supported by SPCImage software, moving average of adjacent pixels). A measured IRF was used in the fit to a biexponential decay model (see below). Regions of interest were identified in FIJI;^16^ lifetime is reported as the average of the weighted average τ_m_ over a region.

### Zeiss LSM 880 with PicoQuant FLIM

Data were acquired on an inverted LSM 880 confocal microsope (Carl Zeiss AG, Oberkochen, Germany) equipped with a FLIM upgrade kit (PicoQuant GmbH, Berlin, Germany). The LSM 880 was controlled with Zen Black software (Zeiss); the TCSPC unit was controlled with SymPhoTime 64 software (PicoQuant). A 485 nm diode laser operating at a repetition rate of 40 MHz was used to provide pulsed excitation for FLIM. Emission was collected through a 40x oil immersion objective immersed in Immersol 518F (Zeiss) and was filtered through an additional bandpass emission filter (550/49 nm, Semrock) before it reached the detector. Single photons were detected with a PMA-Hybrid 40 detector (PicoQuant) and processed with a TH260 Pico Dual TCSPC Unit (T3 TCSPC mode). Data were acquired at an approximate frame rate of 1 Hz; successive frames were binned before analysis to obtain a total acquisition of 5 seconds (5-6 frames per V_mem_ step). A confocal pinhole of ~2-3 Airy units was used to increase photon counts at the expense of optical sectioning.

Fluorescence lifetime analysis was performed with global analysis in SymPhoTime (PicoQuant). All photons from a defined ROI per frame were combined for a global fit (see Appendix 1 for further explanation). An experimentally measured IRF was used for fitting to a biexponential model (see below). Shift was fixed to 0, and offset was determined by the software from the baseline. Lifetimes of successive frames at the same V_mem_ were similar. Data presented here were averaged across all frames at a given potential for each cell before lines of best fit were determined.

### Two-Photon (2P) Fluorescence Lifetime Imaging

2P-FLIM was performed on an inverted Zeiss LSM 510 equipped with a Becker and Hickl SPC-150N photon counting card. Excitation was provided by a MaiTai HP Ti:Sapphire laser tuned to 820 nm with a repetition rate of 80 MHz. Excitation power was controlled by a half-wave plate followed by a polarizing beamsplitter, as well as an acousto-optic modulator (AOM) controlled by the Zen software (Zeiss). Light was directed into the microscope from a series of silver mirrors (Thorlabs or Newport Corporation).

Emitted photons were collected with a 40x/1.3 NA EC Plan-Neofluar oil immersion objective (Zeiss) and detected with an HPM-100-40 single photon counting detector (Becker and Hickl). Photons were detected from the non-descanned side port of the LSM 510 after being reflected off of a 680 nm long pass dichroic mirror and passing through an IR-blocking filter. Emission light was further filtered through a bandpass emission filter (550/49 nm, Semrock). Reference pulses for time correlated single photon timing were sampled from the excitation beam and detected with PHD-400-N high speed photodiode (Becker and Hickl).

2P-FLIM data for VF2.1.Cl were fit to an incomplete biexponential decay model in SPCImage (Becker and Hickl) as described below.^6^ Using the standard solutions described below, the optimal value for the color shift between the IRF and the measured decay was determined to be 0.7 (out of 256 time channels on the analog-to-digital converter). The photon count threshold for fitting was set to 200 counts in the brightest time channel. Other fitting parameters and binning were as described for the published one photon signal.^6^ Lifetime images in lifetime-intensity overlays are scaled as indicated; underlying photon count images are scaled to maximize contrast.

### Measurement of Instrument Response Function (IRF)

For all instruments, the IRF was determined experimentally from a solution of 0.5 mM fluorescein and saturating (12.2 M) NaI in 0.1 N NaOH. The IRF was measured frequently (at a minimum daily) during imaging. The IRF was cropped to only include the time bins with the major pulse (approximately 10 bins in the analog-to-digital converter space). Standard solutions of 2 μM fluorescein in 0.1 NaOH and 1 mg/mL erythrosin B were used to assess quality of lifetime determinations and to refine fit parameters.

### Fluorescence Lifetime Biexponential Fitting

The fluorescence lifetime of VF2.1.Cl was modeled as a biexponential decay (eqn. 1) before convolution with the experimentally measured IRF. Lifetimes are reported as the weighted average τ_m_ (eqn. 2), consistent with the reported analysis for VF-FLIM.^6^

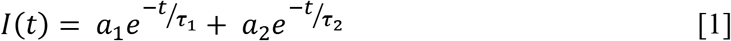

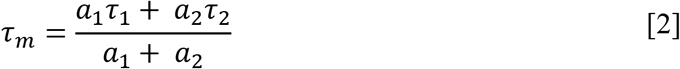

where I is the time resolved fluorescence intensity as a function of time t, a_1_ and a_2_ are the amplitudes of each exponential component, and τ_1_ and τ_2_ are the decay constants (lifetime) of each exponential component. We used both commercial software and custom Matlab code to perform these analyses. The custom Matlab (Mathworks) code is available on GitHub (https://github.com/jlazzaridean/flim_flam/tree/bioelectricity_submission) and documented further in **Appendices 1 and 2**.

We use τ_m_ as a proxy for voltage because it can be more precisely determined with the relatively low photon counts that are typical of biological samples. Further commentary on this is provided in Appendix 1. For calibration of τ_m_ versus V_mem_, we performed a linear fit using Microsoft Excel or Matlab. We quantify the variability measured between calibration samples using intra- and inter-cell variability as described previously.^6^ Briefly, we calculate the root-mean-square deviation (RMSD) between our electrophysiologically determined V_mem_ and the calculated V_mem_ from the lifetime. The “intra” cell variability reports on the error one might expect in successive VF-FLIM measurements on the same cell, and the “inter” cell variability reports on the error one might expect in comparing V_mem_ of two different cells using VF-FLIM.

## Results

### 1P VF-FLIM is Consistent across Different Photon Counting Instruments

To evaluate the portability of the VF-FLIM technique to other laboratories, we developed a standardized measurement scheme for calibrating lifetime with respect to V_mem_. Whole-cell voltage clamp electrophysiology allows us to set V_mem_ of a cell of interest to a defined value. If fluorescence lifetime is measured during these electrophysiological steps, a lifetime-V_mem_ curve can be constructed, as described previously.^6^ We selected this experiment as a comparison point across systems because it is both precise and relatively easy to perform on accessible samples (HEK293T cells).

We measured the lifetime-V_mem_ relationship for VF2.1.Cl on two independent time-correlated single photon counting (TCSPC) instruments (**Figure 1**, **Table 1**). These two TCSPC systems were both distinct from the instrument used in the previously published work.^6^ All instruments in this comparison were both point-scanning confocal instruments with excitation at approximately 488 nm. However, there were considerable differences among the instruments: each was equipped with a distinct microscope bodies, light sources, detectors, and photon counting modules. Furthermore, the analysis was performed in two different commercial software packages and with different binning schemes. Additional instrumentation details are provided above in the Methods.

**Table 1.** Summarized Lifetime-V_mem_ Calibration for VF2.1.Cl in HEK293T on Different TCSPC Systems. Summary of data presented in **Figure 1**. Slope, y-intercept, and intra-cell root-mean-square deviation (RMSD) values were averaged across all cells measured on a particular instrument and are presented as mean ± SEM. Data represent the following numbers of cells: VF-FLIM 17 (data from previously published study),^6^ Zeiss/PicoQuant (PQ) 6, Nikon/Becker-Hickl (BH) 7.

| System | Slope (ps/mV) | Y-Intercept (ps) | Inter-cell RMSD (mV) | Intra-cell RMSD (mV) |
| --- | --- | --- | --- | --- |
| VF-FLIM<sup>6</sup> | $3.50 \pm 0.08$ | $1.77 \pm 0.02$ | 19 | $3.5 \pm 0.4$ |
| Zeiss/PQ | $3.43 \pm 0.08$ | $1.75 \pm 0.03$ | 19 | $3.1 \pm 0.6$ |
| Nikon/BH | $3.18 \pm 0.03$ | $1.80 \pm 0.02$ | 14 | $3.4 \pm 0.5$ |

**Figure 1.**
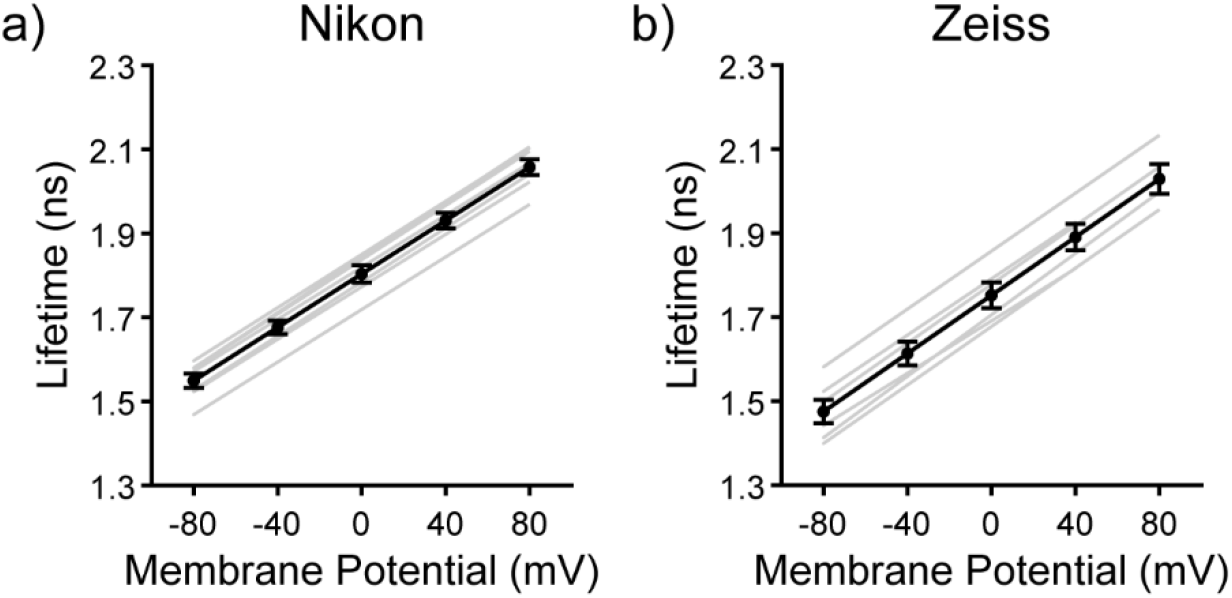
Lifetime-V_mem_ calibrations for VF2.1.Cl on different TCSPC systems. HEK293T cells were held at specified V_mem_ with whole-cell voltage clamp electrophysiology while lifetime was measured on a) a Nikon confocal equipped with a Becker&Hickl FLIM system and b) a Zeiss LSM 880 confocal equipped with a PicoQuant FLIM system. Gray lines indicate lines of best fit for individual cells; black line indicates the average line of best fit. Data are shown as mean ± SEM.

We observed strikingly similar lifetime results across all three systems in this comparison. The slopes, y-intercepts, and spreads among the different calibrations are nearly identical (**Table 1**). Furthermore, the voltage resolution obtained on all three systems is similar. To quantify this, we calculated “intra-cell” and “inter-cell” accuracy in the calibration, as described previously.^6^ Intra-cell variability captures the expected error in successive measurements on the same cell, while inter-cell variability captures the expected error in measurements of resting membrane potential differences between different cells. We find that all three instruments provide similar accuracies in lifetime-based V_mem_ determination. Notably, the intra-cell resolution achieved with VF-FLIM (3.1-3.5 mV) is approximately 6-fold better than what we could achieve using a ratio-based approach with di-8-ANEPPS (18 mV resolution), and approximately 10-fold better than what we could achieve using lifetime measurements with CAESR (33 mV resolution). The inter-cell resolution of VF-FLIM (14-19 mV) also compares favorably (10 to 20 fold improvement) to ratio-based techniques (150 mV resolution) or lifetime with CAESR (370 mV resolution).^6^

### VF2.1.Cl Fluorescence Lifetime Reports Voltage under Two-Photon (2P) Excitation

We next sought to determine whether the fluorescence lifetime of VF2.1.Cl under 2P excitation could be used to make determinations of absolute membrane potential. For these measurements, we used a FLIM instrument similar to the one described previously,^6^ but with an altered beampath to enable 2P excitation (see Methods above). We selected an excitation wavelength of 820 nm, near the previously reported 2P absorption maximum of fluorescein.^17^ Excellent membrane staining and optical sectioning was observed (**Figure 2a**). Due to the relatively poor two photon cross section of VF2.1.Cl,^18^ high light power was required to obtain sufficient photon output. As such, photobleaching appeared more rapid under these conditions than under 1P excitation with similar photon output (data not shown), but it was still possible to obtain high-quality patches.

**Figure 2.**
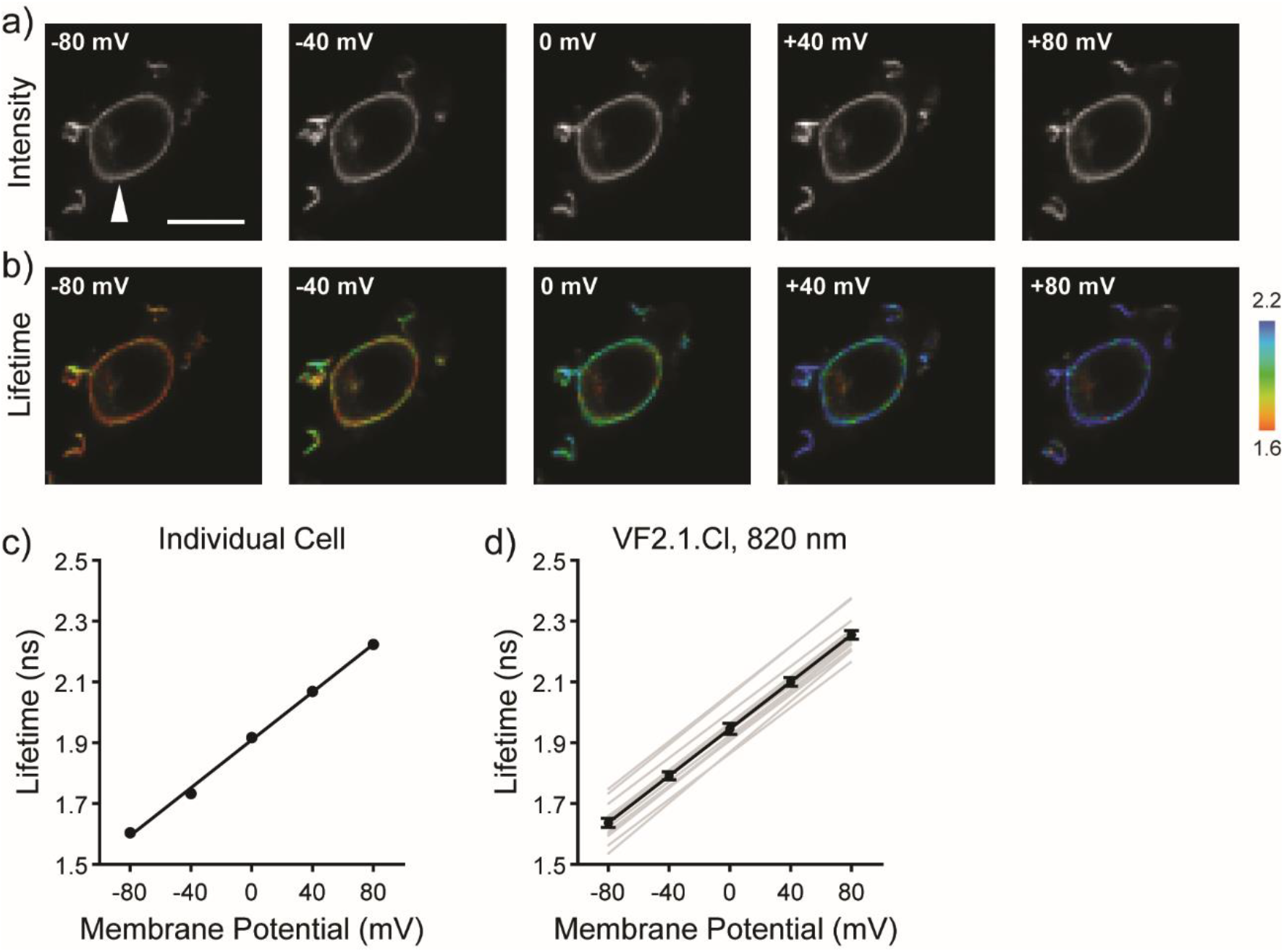
VF2.1.Cl fluorescence lifetime reports absolute V_mem_ under 2P excitation. Simultaneous whole-cell patch-clamp electrophysiology and two photon FLIM (820 nm excitation) of VF2.1.Cl in HEK293T reveals a V_mem_ sensitive τ_fl_. **a)** Photon count images of 100 nM VF2.1.Cl in a HEK293T cell held at the indicated V_mem_. White arrow indicates the patch pipette; scale bar is 20 μm. **b)** Lifetime-intensity overlay of the same HEK293T. **c)** Quantification of the lifetime images in (b), revealing a clear linear relationship between τ_fl_ and V_mem_. Points indicate average lifetime for the individual cell in (b) within a membrane localized region of interest at each potential. Black line indicates the line of best fit. **d)** Aggregated lifetime-V_mem_ calibration for VF2.1.Cl, where individual cells are represented by gray lines. Black line represents the average slope and y-intercept for n=16 HEK293T cells. Data are shown as mean ± SEM.

Using simultaneous fluorescence lifetime imaging and whole cell patch clamp electrophysiology as described above, we determined that the τ_fl_ of VF2.1.Cl under these excitation conditions is sensitive to V_mem_. With 820 nm 2P excitation, we observe a sensitivity of 3.86 ± 0.04 ps/mV with a 0 mV lifetime (y-intercept) of 1.95 ± 0.01 ns (n=16 cells, **Figure 2**). These results are qualitatively similar to the 1P calibrations (**Table 1**), but we do observe consistently larger slopes and y-intercept values under 2P illumination. Using this 2P calibration, we observed similar or slightly improved V_mem_ resolution to the 1P calibration (14 mV inter-cell RMSD and 4.4 ± 0.6 mV intra-cell RMSD for the 2P set-up, calculated as in **Table 1**).^6^

## Discussion

To broaden the reach of the VF-FLIM^6^ technique and standardize its use, we compared the fluorescence lifetime of VF2.1.Cl in HEK293T cells under whole-cell voltage clamp across entirely different TCSPC systems, as well as under 1P and 2P illumination. We find that the results are similar across different lifetime instruments and configurations, consistent with the intrinsic nature of fluorescence lifetime. This observation highlights a key advantage of fluorescence lifetime as opposed to fluorescence intensity, namely its relative independence from illumination intensity and dye concentration-related artifacts. In addition to demonstrating the robustness of VF-FLIM, the expansion of the technique to 2P illumination opens up the possibility of measuring V_mem_ in thick tissue, enabling studies of V_mem_ during more complex developmental processes.

### VF-FLIM is Reproducible across Lifetime Instruments

We compared the results of a standardized electrophysiology experiment (5 voltage steps in HEK293T cells) across three TCSPC FLIM microscopes (1 from published data and 2 additional microscopes). We observe excellent agreement between these systems, suggesting that a calibration obtained on one TCSPC FLIM system could be used on a different system with minimal recalibration. Taken together, our results demonstrate that VF2.1.Cl lifetime is a robust proxy for membrane potential across a variety of TCSPC configurations.

Nevertheless, there were slight differences in calibration measured between the instruments, so the most accurate V_mem_ determinations would require electrode-based calibration on the system to be used for data collection. These slight discrepancies could result from a variety of factors, including differences in temperature, stray light, IRF stability, or simply random variation between cells. We note that the slightly reduced inter-cell RMSD on the Nikon instrument is difficult to interpret. These data were collected over a much shorter time frame than the previously published results (~2 weeks versus 1.5 years), so the effects of any instrument-related drift would be diminished.

Along with these data, we have also provided extensive commentary on the processing of VF-FLIM data (**Appendices 1 and 2**), together with a custom Matlab codebase for data analysis. We have observed that incorrect selection of the fitting model and parameters can introduce sizable artifacts into the lifetime results, often much larger than the differences we observe between microscopes (**Table 1**). We hope that our added notes will facilitate the broader use of VF-FLIM for absolute membrane potential determinations.

Because of limitations in our access to diverse FLIM instrumentation, we have not evaluated the performance of VF-FLIM in a frequency domain (FD) configuration. Exploration FD FLIM would offer the opportunity to make considerably faster lifetime recordings in a widefield imaging configuration. Because many interesting V_mem_ dynamics occur on shorter timescales, investigation of VF-FLIM in combination with fast FD techniques such as siFLIM^19^ are of considerable future interest.

### 2P Absolute Voltage Determinations with VF2.1.Cl

We determined that the fluorescence lifetime of VF2.1.Cl under 2P illumination is sensitive to the transmembrane potential. This observation is a critical step forward in 2P voltage imaging, as translation between 1P and 2P illumination is not always straightforward for V_mem_ indicators. For example, a comparative study of genetically encoded voltage indicators (GEVIs) revealed that many of them did not display sufficient brightness or sensitivity to record V_mem_ under 2P illumination.^11^ However, we were optimistic that the PeT mechanism of the VoltageFluor dye VF2.1.Cl would be translatable to 2P lifetime imaging, based on previous work using VoltageFluors in 2P intensity-based imaging.^18,20^

Although it is not strictly required, fluorophore emission spectra and lifetime are generally similar under 1P and 2P excitation.^21^ Consistent with this, we observe good agreement between the properties of VF2.1.Cl under 1P and 2P illumination. The slight increase in the y-intercept (0 mV lifetime) is most likely attributable to reduced contribution of cellular autofluorescence when exciting at 820 nm (2P) versus 479 nm (1P). This hypothesis is consistent with the slightly increased sensitivity under two photon illumination, as autofluorescence will contribute a V_mem_-insensitive background to the lifetime recording. This reduction in autofluorescence could also produce the observed slight resolution improvement for V_mem_ recordings (inter-cell variability of 14 mV in 2P versus 19 mV in 1P). Nevertheless, these data were collected over the course of a much smaller time window than the 1P data in our previous study (1 month versus 1.5 years). Slight differences in instrument performance over time may also contribute to this 1P versus 2P resolution difference.

Our work demonstrates that the fluorescence lifetime of VF2.1.Cl under 2P excitation could be a useful method for measuring absolute V_mem_, as it displays excellent V_mem_ sensitivity and resolution. However, a key limitation of the work presented here is the poor 2P cross-section of the dichlorofluorescein chromophore in VF2.1.Cl. Expansion of VF-FLIM to additional VoltageFluors and colors,^22^ including those with improved 2P cross-sections,^18,20^ will likely lead to considerable improvement in performance. Together, we hope these technologies can begin to address important questions of V_mem_ signaling over long timescales and in thick tissue.

## Conclusion

We demonstrate that the fluorescence lifetime of the VoltageFluor dye VF2.1.Cl (VF-FLIM) is a reproducible and robust measure of absolute membrane potential across different instruments and illumination modes (1-photon and 2-photon). We also offer extensive documentation of our data processing routines in the hopes that it will facilitate more widespread adoption of the VF-FLIM technique. We are optimistic that the flexibility and robustness of VF-FLIM will enable its application to study the diverse biological roles of membrane potential across space and time.

## Acknowledgements

Research in the Miller lab is supported in part by the NIH (R35GM119855). J.R.L.-D. was supported in part by an NSF Graduate Research Fellowship. We thank Holly Aaron and Feather Ives for expert technical assistance. A subset of FLIM experiments were performed on “Deckard” at the UC Berkeley CRL Molecular Imaging Center, supported in part by NSF DBI-0116016.

## Appendix 1. Technical Notes on Fitting VF-FLIM Data

## Image Analysis: Selection of Global versus Pixelwise Analysis

Fluorescence lifetime analysis in the context of images can be broken down into two major classes: pixelwise and global (**Scheme 1**). In pixelwise analysis, the fluorescence decay constants τ are determined at each pixel and a heatmap of τ is generated. For summaries of results, the average of these τ values in the region of interest is often obtained. Global analysis involves consolidation of photon histograms for an entire region of interest (ROI) *before fitting*; decay constants are then determined per region of interest from the selected exponential model.

**Scheme 1.**
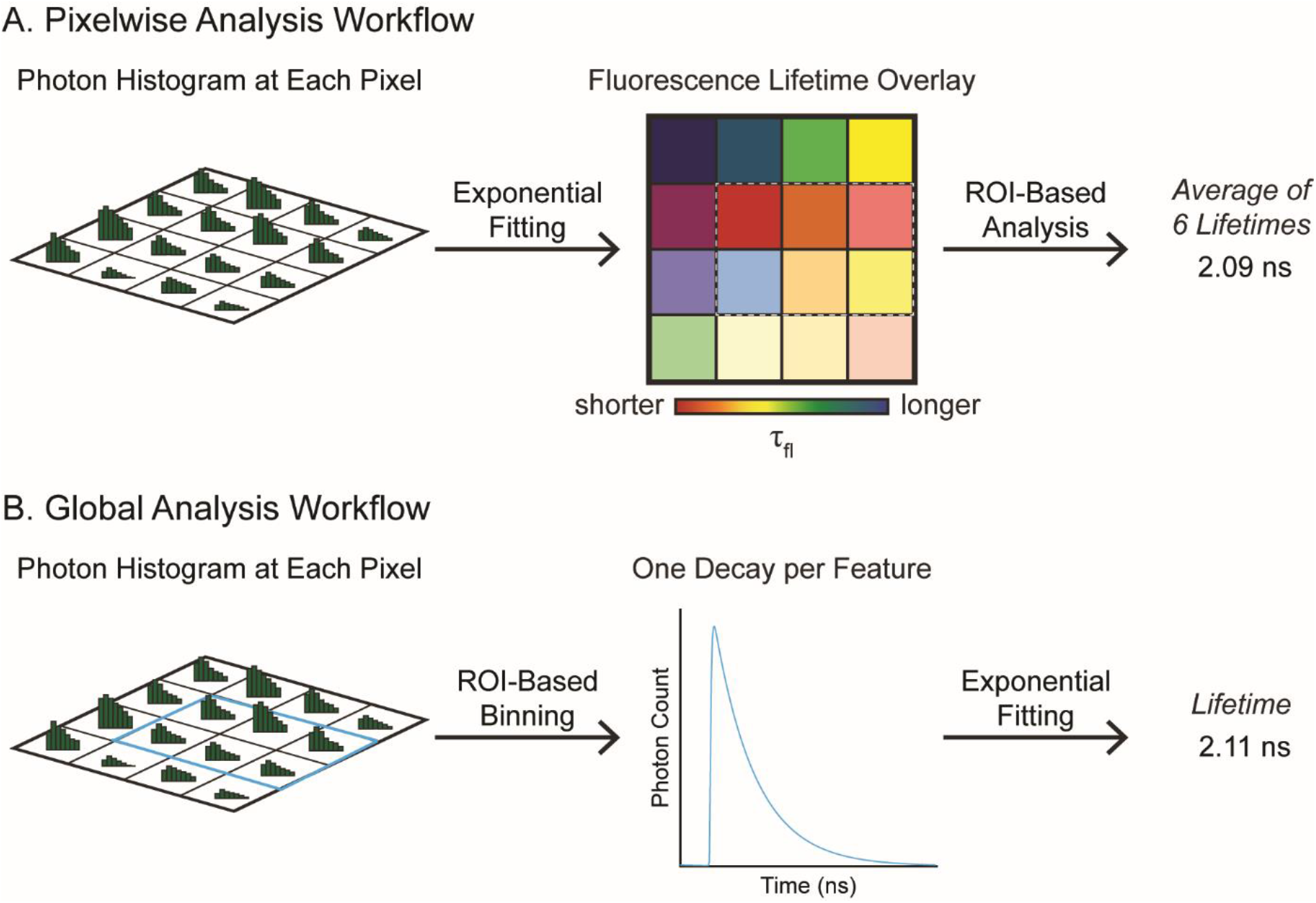
Workflows for analysis of fluorescence decays. (A) Analysis at each pixel involves fitting an exponential decay model to each position in the image. Often, images must be binned dramatically to obtain sufficient photons to perform each fit From this binned image, a region of interest (ROI, dotted line) is identified. The average across the pixels in the ROI is often used to represent the lifetime per ROI. (B) Global analysis involves combination of raw photon histograms from multiple pixels in the ROI (blue outline), followed by fitting of an exponential fit on the combined decay. These two approaches give similar results for large numbers of photons.

The selection of pixelwise versus global analysis methods depends largely on the application at hand, and there is no general rule. We have used both pixelwise and global analysis with VoltageFluor FLIM data. Indeed, an advantage of the attached FLIM-FLAM code package is that it enables facile global and pixelwise analysis on the same dataset. Below, we outline some considerations in selecting an analysis mode.

On the one hand, in using VF-FLIM to map V_mem_ in a biological specimen, pixelwise analysis is often the most visually satisfying. Pixelwise analysis also provides an easier interface to discover unexpected relationships in the data, as the regions with interesting features do not need to be known a priori. However, care must be taken in the generation of images fit at each pixel, as many photons are required to accurately fit exponential decays. In many cases, acquiring enough photons to resolve the lifetime at each pixel limits the temporal resolution of the experiment. Most pixel by pixel fitting with VoltageFluors shows only the weighted average decay, as the individual decay constants and their amplitudes are considerably noisier.

If detailed information is desired about individual components of multiexponential fluorescence decays, global analysis is usually the best option. Sufficient photons to determine a fluorescence decay can be obtained much more easily on sensitive specimens, and often enough photons may be collected to resolve individual parameters of the decay model. Global analysis can also be used to extract information from carefully constructed regions of interest on the raw photon image, which may be useful for identifying fluorescence decays from fine structures such as neuronal processes.

## Selection of an Exponential Decay Model

The simplest fluorescence decay model is a single exponential decay, reflecting a uniform population of molecules emitting with a decay constant τ. In practice, many fluorescence decays are not well described by a single fluorescence decay model. Multiple populations of emitters exist in most biological samples, resulting from heterogeneous probe environments and the fluorescence of endogenous chromophores. These multiple populations of emitters result in a fluorescence intensity I that decays over time t as a sum of exponential decays, each with amplitude ai and decay constant τ_i_ (eqn. A1-1).

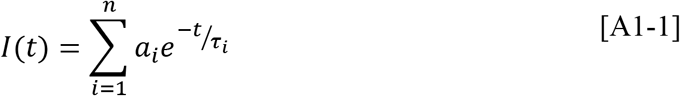

Selection of the most appropriate number of decay components is a statistically challenging problem. With VoltageFluors, a naïve expectation would be that the decays would show a single exponential form, arising from fluorophore uniformly localized to the interface between the plasma membrane and the extracellular space. However, in most cases, VF2.1.Cl in cells is best described by a biexponential fluorescence decay.

For fluorophores delivered to biological systems, a practical decay model is often selected by iteratively fitting additional terms and monitoring the concomitant decrease in the reduced chi squared. We demonstrated this approach when we reported a new suite of VoltageFluor dyes with FLIM characterization.^23^ This model selection process is best performed on a standardized dataset. Ideally, the standards would be close to the system of interest (e.g. voltage-clamped HEK293T cells to develop standard fitting for non-voltage clamped HEK293T cells), but if that is not possible, lifetime standards characterized by other researchers can be used.^24^ The addition of terms must also be appropriate for the total number of photons acquired, as overfitting can produce artifacts in the results.

## Selection of Other Fit Parameters

After the exponential model is selected, the most important additional parameter in the model is the shift in time between the instrument response function (IRF) and the fluorescence decay (sometimes called the color shift), as well as whether this parameter is fixed or allowed to vary during fitting. We follow the common convention of working with shift in units of bins in the photon histogram. In the data presented here, 1 bin is approximately 0.05 ns. A measured IRF at the same wavelength as the sample should have a shift very close to 0, but a measured IRF at a different wavelength may show a shift in time, especially on older detectors. In our experience, the best way to optimize the shift is to fit a monoexponential lifetime standard with the same emission spectrum as your fluorophore and then fix the shift to the shift obtained for that standard. Synthetic IRFs determined from the rising edge of the fluorescence decay are not implemented in FLIM-FLAM. Although they are common in commercial FLIM packages, we found a dramatic increase in the noise in VoltageFluor lifetimes determined from calculated IRFs, likely attributable to poor modeling of the short τ components present in certain VoltageFluor fluorescence decays.

The offset of the lifetime decay is the amount of non-time-resolved signal in the decay, arising most commonly from dark counts on the photon counting detector or from stray light in the room. If acquisitions were carefully performed in a cool, dark room, this parameter can generally be fixed to zero. When the lifetime of the fluorophore is much shorter than the period in between laser pulses, the offset can also be directly measured from the flat baseline of the TCSPC data.

The start and end time bin of the IRF should be selected to include the main IRF pulse but avoid other noise in the baseline. The start and end time of the decay should be a few bins in from either end of the signal region to avoid artifacts from the time to amplitude converter (TAC). Using the average of two measured IRFs in lieu of single, raw measured IRF can correct for variability in any individual IRF but is generally not required. The threshold in number of photons for fitting a decay depends on the number of parameters in your model; more photons are required to model more decay parameters. Whether enough photons have been counted is usually determined empirically based on the consistency of the result and the observation that collecting additional photons does not change the resulting lifetime. For VF2.1.Cl, we generally work with a minimum of 5000 photons total per decay trace, but the consistency of the lifetime obtained at a given number of photons should be checked for each new system, probe, and fit model.

## Appendix 2. “FLIM-FLAM” Exponential Fitting Code User Manual

Time-correlated single photon counting (TCSPC) data are generally fit to exponential decay models, allowing results to be represented with summary parameters such as the weighted average lifetime. While both commercially available and open source software for this purpose exist, we found that no individual package implemented all or even most of our required functionalities. To remedy this, we developed a MATLAB pipeline (FLIM fluorescence lifetime analysis module, or FLIM-FLAM) capable of processing fluorescence lifetime imaging data in an automated fashion. This improvement to the analysis workflow was essential both for development of reproducible fitting models and for the advancements in throughput made by VF-FLIM.

The purpose of FLIM-FLAM is to process, analyze and visualize fluorescence lifetime imaging data. The specifics of this workflow were optimized towards membrane-localized fluorescence signal in mammalian cultured cell lines, but much of the package is generalizable to any application.

This workflow is broken down into three conceptual steps (below). Our modular design allows each of these steps to be called individually or as part of a sequential workflow.

1. Process: Identify the regions of interest (or pixels) within TCSPC data. Extract decays for fitting.
2. Analyze: Fit the fluorescence decays to the specified model. In this case, the model is a sum of 1, 2, 3, or 4 exponential decays.
3. Visualize: Rebuild the fluorescence lifetime data into image format or summarize it by time point and region of interest for plotting of summarized results.

The fitting is performed through iterative reconvolution of a guess model with the instrument response function (IRF). Here, we use a weighted least squares algorithm, together with the built-in fmincon optimization algorithm in Matlab. In addition to the fit output, the program will provide various estimates of goodness-of-fit, as well as inputs re-saved in .mat format for easy reprocessing. Most output files will by default save to the folder where the input data are stored.

The code requires raw photon counting histograms (.sdt or .bin exports), as well as metadata and configuration files. See below (and included examples) for documentation of the format of these additional files. The code requires installation of the BioFormats Matlab package for parsing the .sdt file format for raw photon count data from Becker & Hickl TCSPC systems. Additional documentation can be found within the code files and working examples.

## Data Import

The wrapper code for the file parsers (importData.m) accepts four different import modes, which determine how the data are broken down. The below descriptions depict how an image of X pixels by Y pixels by N time channels would be handled by the software.

1. “single”: Combines all of the pixels X by Y into a single N by 1 decay for each image.
2. “pxwise”: Retains the native spatial resolution and returns an X by Y by N array.
3. “global_fiji”: Parses ROIs provided by the user as binary exports from ImageJ (FIJI). This mode uses these masks to define regions of interest for global analysis.
4. “global_btm”: Uses thresholding code optimized for finding membranes in confocal images of cultured cells to identify ROIs for global analysis. The user has an opportunity to modify and verify ROIs generated by this “batch trace membranes” (BTM) routine.

Once the data have been imported, there are two exponential fitting functions easily available to the user. For global analysis, fitDataStruct.m will process the output of a global import and return a structure with the exponential decay fit results as well as any metadata. For pixelwise analysis, pxwiseAnalysis.m will fit the output of a “pxwise” import and render images of the components of interest, as well as overlays of these components on the photon count image. Average lifetimes for regions of interest in the images can be determined after pixelwise fitting using the provided script batchTraceFromStruct.m. This implements a similar routine to the segmentation performed in global_btm but acts on the fit images instead of the raw photon count histograms.

## Region of Interest Identification

The scripts adapt the routines published previously in batchTraceMembranes2^6^ to enable automated identification of contiguous membrane regions. These regions can be applied either before the fitting (global analysis) or after the fitting (pixelwise analysis). The user has the opportunity to remove debris, merge ROIs that were erroneously split, and approve the final ROIs within the batch tracing script. This processing mode works best if the thresholding is tweaked slightly to reflect the morphology and characteristics of the cell line, so the function takes the cell type as an input parameter. Currently supported cell lines are A431, CHO, HEK293T, MCF-7, and MDA-MB-231. In addition, the option HEK293T_red is available for settings with reduced autofluorescence; use this if HEK293T is causing oversegmentation. If the cell line name is not recognized, the defaults for HEK293T are used.

If no spatial resolution is desired, the global decay for the entire photon histogram can also be selected. More complex ROIs should be generated as TIFF masks (e.g. in FIJI). These can then be imported and applied as masks to the data (0 is background and values >1 are kept).

## Function Usage Overview

The sample sequences of function calls (all MATLAB code files) for five common workflows are listed here for reference. Additional documentation on usage is available within each individual function file (*.m).

1. Pixelwise fitting and rendering of FLIM images: importData, pxwiseAnalysis
2. Pixelwise fitting of FLIM images, followed by averaging across automatically identified membrane regions of interest: importData, pxwiseAnalysis, batchTraceFromStruct
3. Global fitting, in any mode (single decay per image or decays defined by regions of interest): importData, fitDataStruct
4. Rendering TIFF photon count images for defining ROIs on data without processing in any other way: renderPhotonsImg
5. Rendering of TIFF images of parameters that were not previously processed from a pixelwise analysis: renderImage

## Formatting of Input FLIM Data Files and Metadata Files

## Filename Formats for Data Records

FLIM records should be imported as photon histograms generated either by the binary export in SymPhoTime (*.bin) or as the raw acquisition format in SPCM (*.sdt). Opening of sdt files relies on the BioFormats package for MATLAB. Data records can be uniquely identified by a combination of the following parameters:

1. Date recorded, generally YYYY-MM-DD
2. Coverslip or sample identifier (abbreviated ‘cID’, CC).
3. Image identifier (abbreviated as ‘imageID’, II). This is a unique field of view within the sample.
4. Replicate identifier (abbreviated ‘repID’, RR). This is the number of times a FLIM images was recorded from a particular field of view. It is almost always 1, but this parameter is included to allow for re-acquisition of images if technical difficulties arose or for comparisons of signal stability.
5. Frame identifier (abbreviated ‘frameID’, XX). This is the slice of a time series (in experiment time) that the image corresponds to. It is more generally used than replicate ID. For instance, if I recorded two time series of ten images each from the same field of view, they would have the (repID, frameID) combinations of (1, 1-10) and (2, 1-10).

We save our FLIM files to a regularized format so that the file name strings can be parsed to obtain information about the various IDs in the recordings. If filenames are not in the format below, they will need to be renamed for proper functioning of the multilevel ID system.

General format of data filename string, before the .bin or .sdt file extension:

YYYY-MM-DD_CC-II-RR_anything_else_you_wantXXX

**Table A1-1:** Sample parsing results for various filename strings. This is not an exhaustive list of acceptable strings, but it is intended to give the user a sense of parser requirements. The software is flexible regarding the number of digits in CC, II, and RR as long as they are separated by dashes. 02-03-02 and 2-3-2 will give the same result. The RR is frequently not present; it will be assigned to 1 if it is missing. The XXX frameID is only processed if nFrames is set in the metadata (see below). The character before the start of the frameID does not matter as long as it is not another number. Not applicable (N/A) means that the field will not exist in the output data.

|  | Parsing Results |  |  |  |  |
| --- | --- | --- | --- | --- | --- |
| Input name | Date | cID | imageID | repID | frameID |
| 2019-10-12_01-1_bob.bin | 2019-10-12 | 1 | 1 | 1 | N/A |
| 2019-10-12_5-06-3_awesome.bin | 2019-10-12 | 5 | 6 | 3 | N/A |
| 2019-10-12_5-06-3_ralph011.bin | 2019-10-12 | 5 | 6 | 3 | 11 |
| 2019-10-12_010-02_ralph_1.bin | 2019-10-12 | 10 | 2 | 1 | 1 |

## Configuration and Metadata File Formats

This code requires configuration and metadata files to select fit parameters and group data files into categories. In general, these files should be saved in a comma separated values (csv) format in which the first row is the categories and each subsequent row corresponds to a unique record. The analysis suite looks for exact matches for certain categories/fields in the input metadata to properly complete the analysis. Samples of these files are available within the software package, and the fields required are enumerated here:

The <u>configuration file</u> indicates standard fit settings and instrument properties. Most, if not all, of the metadata about acquisition settings is lost upon exporting to a histogrammed photon images (especially in the SymPhoTime software package from PicoQuant), so it needs to be re-entered here. Expected fields are below; order of the columns in the input file does not matter.

1. model: string or character array for the number of exponential components. Accepted values are 1exp, 2exp, 3exp, and 4exp.
2. fixedParam_X: A boolean indicating which parameters (taus and term weights) should be fixed to their starting values (fixedParam_X = 0) and which parameters should be allowed to vary (fixedParam_X = 1) during the fitting. The number _X at the end of the name specifies the parameter, with all of the coefficients numbered before all of the lifetimes. For example, for a biexponential fit, X = [1 2 3 4] for [a1 a2 tau1 tau2]. For a monoexponential fit, fixedParam is set to −1 by default in the software.
3. startParam_X: Floating point fields determining the starting value for each parameter, numbered as for fixedParam_X. If the parameter is fixed, it will be fixed to the input value of startParam_X. floating-point fields for each coefficient and tau to be fit. X is the index of the parameter. For a monoexponential fit, only provide the starting lifetime as startParam_1.
4. offsetMode: parameter indicating whether the offset (dark counts) should be fixed to zero or allowed to vary in the fitting. If offsetMode is 0, then the offset will be allowed to vary in the fitting. If offsetMode is 1, the offset will be fixed to 0. If offsetMode is 2, the offset will be set from the average of the time bins before the start of fitting. If the decay does not completely return to background levels within the period of the laser, offsetMode = 1 should be used (with the caveat that dark counts and stray light background must be low).
5. cShift: floating-point number corresponding to the shift between the IRF and the decay to be fit, in units of ADC time bins.
6. shiftFixed: boolean indicating whether the shift should be fixed to the starting value or used as a free parameter in the fit.
7. nsPeriod: The period of the laser cycle during the acquisition in nanoseconds. For 80 MHz laser repetition rate, the nsPeriod is 12.5.
8. irfStart: The beginning of the IRF (in ADC time bins). This enables cropping of a measured IRF to just the segment corresponding to the laser pulse.
9. irfEnd: The end of the IRF (in ADC time bins).
10. fitStart: The first ADC time bin to be used in the fit.
11. fitEnd: The final ADC time bin to be used in the fit. Note that the fit may stop before this point if a zero is encountered in an ADC time bin to be fit. If zeros are present in the time channels, the fit will stop at the first time bin with zero recorded photon counts.
12. threshold: number of photons minimum in a decay to be processed (sum across all time, not the peak value). This value is only used in pixelwise analysis; global analysis will be performed regardless of the number of photons.
13. irfMode: Accepted values are “single” or “paired.” This parameter indicates whether each IRF file will correspond to an IRF index or if the average of sequential IRF files should be used to generate an IRF for each IRF index.
14. adcRes: The expected number of time channels from the analog to digital converter (N in X by Y by N photon histograms). This value should be constant across all data and IRFs analyzed. It is provided as a fail-safe check in the configuration file to verify that nothing strange happened to the data upon exporting.
15. adcBin: The binning in the time dimension. For example, an adcBin of 1 does not bin, whereas an adcBin of 2 would sum paired adjacent time windows of a 256 bin decay into 128 resulting time windows.
16. viewDecay: Default value of 0 does not show the fit curve for each exponential decay. When set to 1, it will display the fit and residuals for each curve being processed during the fitting processing.
17. maxFunEval: Constraint on the number of function evaluations the fmincon fitting algorithm can perform. See Matlab documentation for more information.
18. stepTol: Convergence criterion in the Matlab built-in fmincon algorithm. See Matlab documentation for more information.
19. optimTol: Convergence criterion in the Matlab built-in fmincon algorithm. See Matlab documentation for more information.
20. constraintTol: Criterion in the Matlab built-in fmincon algorithm regarding bounds. Satisfying it does not stop the solver. See Matlab documentation for more information.
21. ccvColor: Color map for displaying the “color coded value,” usually the lifetime. Accepts Matlab build-in maps (e.g. ‘parula’ or ‘jet’).
22. spatialBin: The bin factor for pixelwise fitting; see binType below.
23. binType: Type of binning to be used with the spatialBin parameter. Acceptable values are (1) ‘std’: square binning (e.g. a *spatialBin* of 2 will lead to 2×2 square binning) or (2) ‘bh’: moving average binning, where each pixel is summed with all pixels *spatialBin* units away, thereby upsampling photons and nominally retaining the native spatial resolution of the image.

The metadata file provides information about the data being analyzed. It is intended to be a useful way to track categorical data and later bring output data with this information into any plotting software. Each file (defined by cID-imageID-repID) must map to a single, unique value in the metadata. The required files are below; again, order of the columns does not matter:

1. irf_index: The number of the IRF to be used in processing the data. If IRFs are imported as an average of a pair of two, this should count the pairs as a group, not as two individual files.
2. cID: The coverslip or sample ID of the record. If no other IDs are provided, all images

Certain special fields in the metadata file are not required, but their presence will change how the data are processed. Any fields beyond the required and special fields will be carried forward and assigned to the appropriate records, but they will not affect the analysis. Special fields are below:

1. imageID: If an imageID is provided, then all imageIDs in the input data must have a match in the metadata, and records will be matched to metadata based on both cID and imageID.
2. repID: Similar to imageID. It is only used for matching records with metadata if it is provided.
3. nFrames: The number of frames in the record. If this is specified, the file name parser will look for and assign the frameID based on the end of the filename.
4. cellType: This field must be provided if the user would like to access the automated membrane tracing in “BTM” modes either before or after fitting exponential decays. Supported values are “A431”, “CHO”, “HEK293T”, “MCF-7” and “MDA-MB-231,” which threshold the image slightly differently to best capture membrane given the typical morphology of that cell line. If cellType does not match one of these values, the code will use the settings for HEK293T cells.
5. smallestObject: This field must be provided if the user would like to access the automated membrane tracing in “BTM.” The units are in pixel areas. The script will assume that regions smaller than smallestObject are debris and will remove them automatically.

If the user is importing ROIs as TIFF masks (e.g. those generated in FIJI, see Materials and Methods), there are two options for matching the ROIs to the relevant images. This option is determined by a command prompt to the user, and it is specified within the function as the mode string (either “csv” or “parse”).

1. parse: Use the structure of the ROI name to determine which image the apply the ROI to. The ROI must be named in the format CC-II-RR_NN, where NN is the numerical identifier of that ROI (the number of ROIs for that image). CC, II, and RR are the cID, imageID, and repID as described above. Names in the format CC-II, CC-II-RR_NN, and CC-II_NN also are accepted. Missing fields are assumed to be equal to 1. Parse mode can be used for time series but the same ROIs will be applied to every frame of the time series. If changing ROIs are desired for a time series, they must be specified in the csv mode.
2. csv: Uses a separate ROI metadata file (*.csv) that contains the cID, imageID, repID, and frameID of the image as well as the exact name of the ROI (minus the file extension). cID and imageID must be present; if repID or frameID is missing, it will be filled in with the value 1.

